# Reliable and automatic epilepsy classification with affordable, consumer-grade electroencephalography in rural sub-Saharan Africa

**DOI:** 10.1101/324954

**Authors:** Vincent T. van Hees, Eric van Diessen, Michel R.T. Sinke, Jan W. Buitenhuis, Frank van der Maas, Lars Ridder, Willem M. Otte

## Abstract

Epilepsy is largely under-diagnosed in low-income and middle-income countries, due to lack of medical specialists and expensive electroencephalography (EEG) hardware. In this study we investigate if low-cost consumer-grade EEG in combination with machine learning techniques can offer a reliable screening tool to improve diagnosis rates.

We acquired brain signals in people with epilepsy (N=163) and healthy controls (N=138) in two difficult-to-reach areas in rural Guinea-Bissau and Nigeria. Five minutes of fourteen channel resting-state EEG data were acquired with a portable, low-cost consumer-grade EEG recording headset. EEG channel time-series were divided in four-second artifact-free epochs and transformed into delta, theta, alpha, beta and gamma wavelet frequencies. Summary measures such as the mean, standard deviation, minimal value and maximal value of the epoch signal fluctuations were used to train a random forest classifier. Epilepsy diagnosis based on at least three months seizure calendar data was used as the gold standard diagnosis. To prevent too optimistic classification the trained model was evaluated with EEG data from subjects not used in the training. In addition, we tested a classification model trained on Nigeria data against data from people in Guinea-Bissau and vice versa. The most contributing data features in the EEG were found in the beta and theta frequencies in Guinea-Bissau and Nigeria, respectively. Within-country model performance was good with area under the receiver-operating curves of 0.85 and 0.78 (± 0.02 standard errors) in unseen data in Guinea-Bissau and Nigeria, respectively. Across-country performance was moderate (0.62 and 0.64 ± 0.02).

Our data suggests that a combination of low cost electroencephalography and machine learning techniques may facilitate diagnostic screening for epilepsy in the most remote areas of the world.

## 1. Introduction

Epilepsy is a severe neurological condition with a prevalence of 65 million cases worldwide. The reported incidence is 45 and 82 per 100,000 per year for high-income and low-middle-income countries, respectively [1]. Even more concerning is that for many low-income and middle-income countries no incidence rate is reported, and over 90% of all people who suffer from epilepsy in low-middle-income countries remain undiagnosed and hence untreated [1]. This is known as the epilepsy diagnostic gap.

The epilepsy diagnostic gap can be explained by the limited diagnostic tools and health care resources in these countries. In many of these places acquisition of clinical-grade electroencephalography (EEG) is not available and the clinical experts to interpret the data are few and mostly concentrated in the larger cities [2]. Furthermore, the interpretation of EEG recordings is a challenge due to their non-stationary nature [3,4], the high inter-observer variation in clinical scoring, and the dependency on trained clinicians [5]. Therefore, automatic ’expert-free’ robust methods to perform an initial selection of individuals with high epilepsy probability would be very useful to optimally spend the limited resources and support community-services.

In this study we explored the potential of affordable diagnostics that circumvent specialist-dependency, long recordings in clinical settings. To that aim we collaborated with two epilepsy services from community-based rehabilitation (CBR) programs in rural sub-Saharan Africa [6]. At least three months of seizure data were required prior to examination and reliable epilepsy diagnosis. A definite diagnosis is based on follow-up data from similar calendars for at least a year.

We investigate whether this time-consuming diagnostic procedure could be replaced by means of several minutes functional resting-state brain data acquisition with a portable, low-cost consumer-grade electroencephalography-recording Emotiv EPOC headset (around $600) [7–10]. This fourteen-channel headset has been introduced by the gaming industry to capture conscious thought, emotions, facial expressions and head rotations and is comparable in signal quality to clinical acquisitions [10].

We hypothesized that the EEG data of people with epilepsy, even in resting-state periods with no seizures, would have a deviating ‘signature’ (in comparison to controls) that could potentially be picked up by combining time-series features and machine learning (see Figure 1 for Infograph). We also hypothesized that these resting-state EEG deviations allow for automatic labeling of presence or absence of epilepsy with sufficient quality to function as a potential screening tool in resource-poor and difficult-to-reach areas.

**Figure 1:**
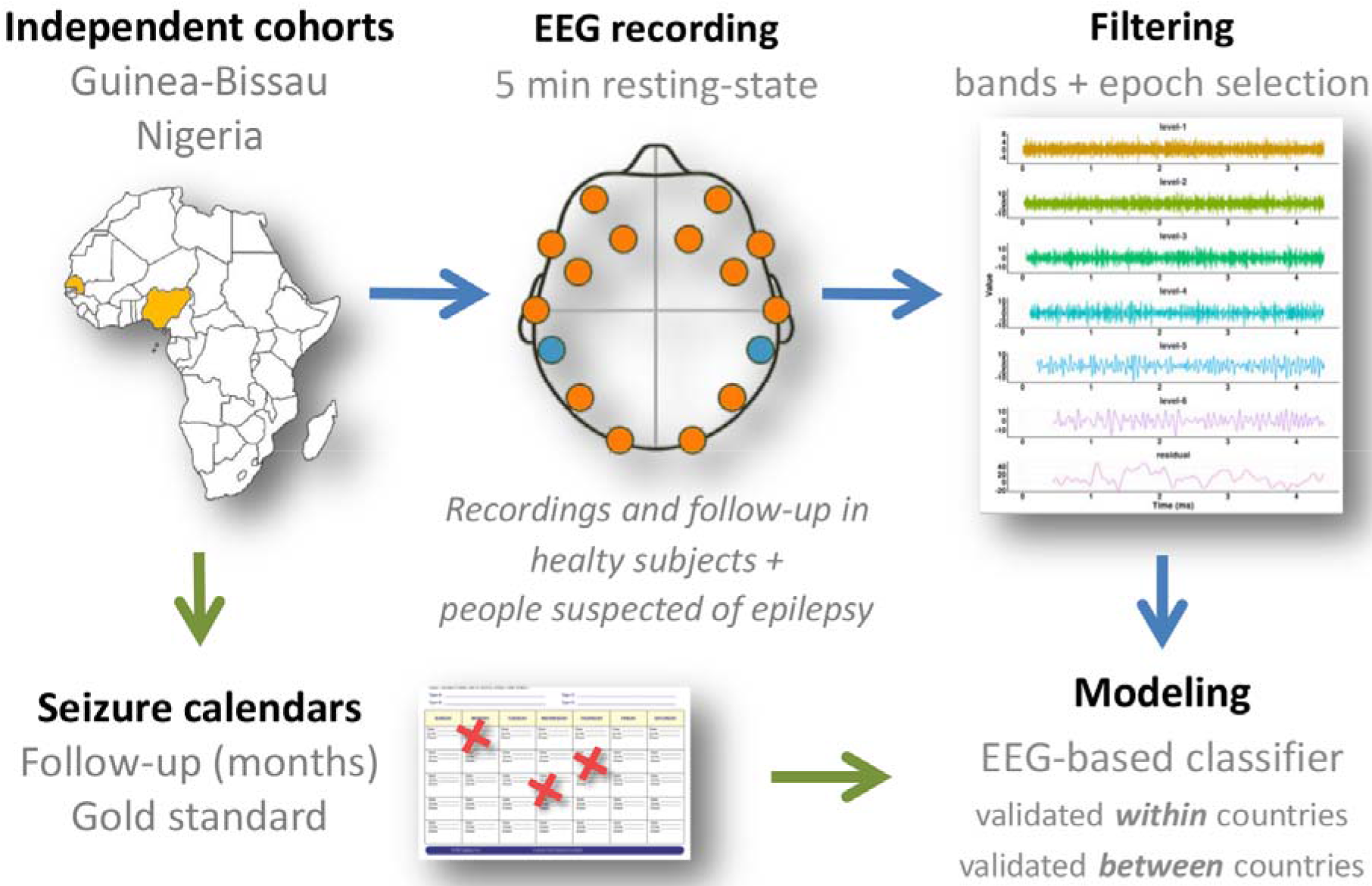
Infograph of the study flow. Two independent cohorts of people are included from community based rehabilitation centers in rural Guinea-Bissau and Nigeria with resting-state EEG recordings in all subjects. People are followed over time. The definite epilepsy diagnosis in the people suspected of epilepsy is determined by means of seizure calendars during follow-up and used as gold standard data for model training. EEG recordings are filtered at channel level in different frequency bands and devided in multiple epochs. Epochs are characterized by simple features: the mean, minimum, maximum and standard devation of the signal. EEG classifiers, which allow prediction of presence or absence of epilepsy, are constructed from these features and validated with unseen data within countries and across countries.

## 2 Materials and Methods

### 2.1 Location

Our study was conducted at a community-based rehabilitation center in Izzi Local Government Area, Ebonyi State, Nigeria and a similar community center in Bissorã, Oio, Guinea-Bissau. These communities aim to reduce the burden of disabilities, including neurodisabilities such as epilepsy. All individuals included in the study lived in the surrounding villages.

### 2.2 Selection of people with epilepsy

Community-based rehabilitation (CBR) programs were used to identify individuals with epilepsy in difficult-to-reach villages. The identification was performed by CBR workers according to a three-step approach, including 1) a general screening of the population, 2) registering of daily seizures on a calendar for at least three months by the individual or a caretaker and 3) the clinical evaluation of the people with epilepsy and the description of the seizures at the rehabilitation center [11]. This approach was recently validated in one of the CBR centers [6].

### 2.3 Ethics

The organizational boards and the local (Secretário de Saúde Bissorã [05–16]) and the national government (Ministério da Saúde, Bissau) approved the study in Guinea-Bissau. The local health ministry (Izzi, Local Government Area) and the federal government (Ebonyi State House of Assembly, Abakaliki [7-11-2016]) approved the study in Nigeria. Informed consent was obtained from adult participants and caretakers of children below eighteen years. Control participants were recruited from healthy caretakers and family members.

### 2.4 EEG acquisition

EEG data was collected in May/June 2016 in Guinea-Bissau and in September/October 2016 in Nigeria with the same headsets (EMOTIV Inc, San Francisco, USA). The fourteen channel EEG was configured to sample at 128 Hertz with a 16-bit resolution. Participants were asked to sit on a chair for five minutes while wearing the wireless headset. Two sensors were preserved for reference and grounding: the ‘common mode sense’ (CMS; located at P3) sensor was used as the active reference for absolute referencing. The ’driven right leg’ (DRL; located at P4) sensor was used for feedback noise cancelation. The electrodes were located at anterofrontal (AF3, AF4, F3, F4, F7, F8), frontocentral (FC5, FC6), occipital (O1, O2), parietal (P7, P8) and temporal sites (T7, T8), according to the International 10-20 system. Signal quality scores are recorded for each electrode with a range of one to five (no units), with five as best quality. Two minutes of resting-state EEG data were recorded with closed eyes during these five minutes and used for our study. The researcher kept a log on deviations from the experimental protocol or unusual events that may affect the experiment. Data was stored on a Bluetooth-connected laptop.

### 2.5 Data cleaning and epoch selection

The data processing is summarized in Figure 2. Visual inspection of the raw signals indicated random temporal jumps in the signal offset in a subset of the datasets. We considered these offsets to be non-physiological and corrected them with a basic interpolation and outlier-removal filter. We corrected for offset drifts with a four second rolling median filter.

**Figure 2:**
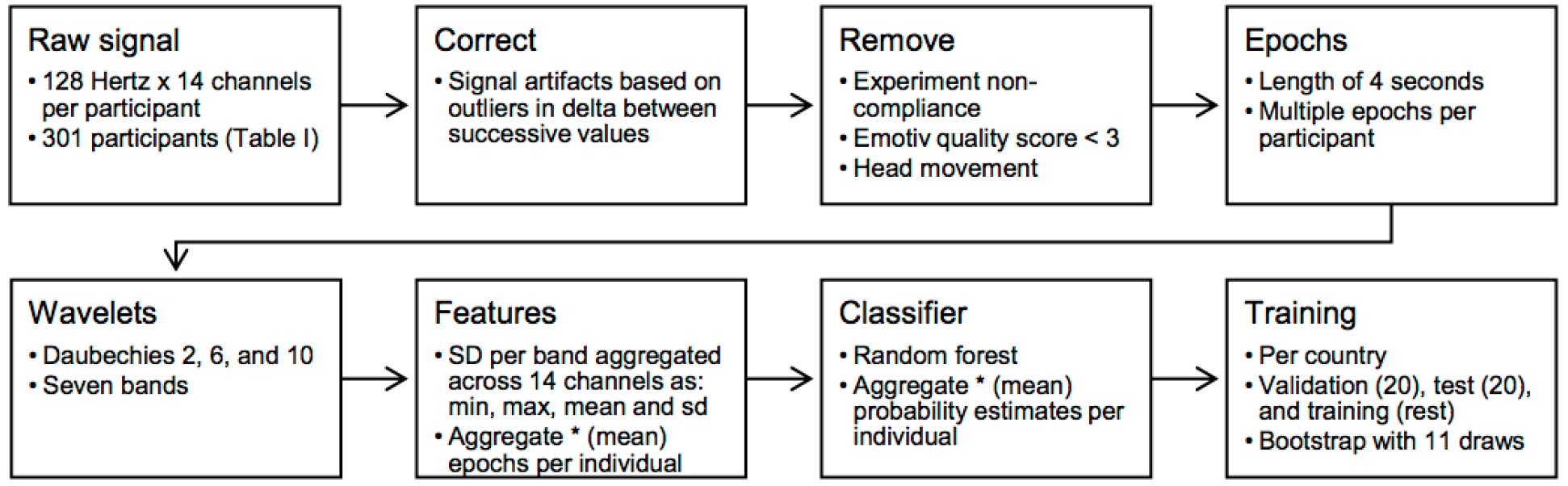
Data processing stages. SD: standard deviation, min: minimum, max: maximum, *: either of these aggregation steps is done, not both.

Next, time segments of the EEG data were removed i) if the research log indicated a deviation from the protocol, ii) if the signal quality score produced by the Emotiv device was 1 or 2 (which are labeled by Emotiv as ‘very bad’ and ‘bad’, respectively) for any of the EEG channels, iii) or if the absolute deviation of the gyroscope signals relative to its overall median was larger than thirty. The gyroscope value threshold of thirty was based on visual inspection of the data, which indicated that most head movements result in a change of more than thirty. All remaining time segments were split into four-second epochs for further wavelet analysis.

### 2.6 Wavelets

In line with previous studies we used multi-resolution wavelet analyses to distinguish frequency bands in the signal [12–14]. Three wavelet types, Daubechies 2, 6 and 10, were derived to cover the commonly used range of Daubechies d1 to d10. Next, seven wavelet levels were extracted per wavelet type: delta (0.5-1 Hz), delta (1-2 Hz), delta (2-4 Hz), theta (4-8 Hz), alpha (8-16 Hz), beta (16-32 Hz) and gamma (32-64 Hz).

### 2.7 Features

Features were based on the fluctuations in the epoch signal. These fluctuations are in indicator of the amount of neural activity in the tissue underlying the EEG channel. We captured these fluctuations with the standard deviation. This standard deviation was calculated for each combination of wavelet type and wavelet level. Next, standard deviations were aggregated across the fourteen channels as minimum, maximum, mean, and standard deviation. With this aggregation across the channels, we expect to reduce the sensitivity of the resulting model to specific localization of epileptic activity, which may vary between people with epilepsy. The aggregation may also make the model less dependent on headset and sensor positioning.

The analyses were performed by either using the features, corresponding to the multiple epochs within an individual, as input for the classification model or by first aggregating the features (mean) per individual before feeding them into the classification model. If no aggregation was used at this point then the classification model will give one probability of epilepsy per epoch, which are then aggregated (mean) per individual (Figure 2).

In summary, from each person with epilepsy all available epochs were used. Per epoch of four seconds and for each of the three wavelet types, 28 features were extracted: the four aggregation types applied to the standard deviations in wavelet values times seven wavelet levels. Next, these features were either aggregated per individual before or after the classification stage (Figure 2).

Our R code and anonymized EEG data can be found online [15].

### 2.8 Classification models

Random forest classification models were trained using supervised learning with the R packages randomForest (version 4.6-12) and caret (version 6.0-73). A Random forest model is an ensemble of decisions trees, which involves making successive binary decisions via the branches of the tree based on feature values. The advantage of using many decisions trees, being a forest, is that it can use more input variables and it lowers the risk of overfitting towards training data [16].

The classification model outcome was binary: epochs originating from ‘epilepsy’ or ‘control’ brains. We used the default model size of 500 decision trees. Here, the number of candidate variables per tree was tuned by random search, using three draws. Features per split were selected with the Gini index, representing the node purity or importance. Finally, accuracy was used as performance metric in five-fold cross-validation. No information about the participants other than the EEG recording was used in the classification process.

### 2.9 Classifier training and evaluation

Models were trained separately for the Guinea-Bissau and Nigeria study population. Models were trained and validated with each of the d2, d6 and d10 wavelet type, leading to a total of twelve Random forest models (2 countries × 2 aggregation options × 3 wavelet types). Here, participants were randomly allocated to a validation set (ten participants with epilepsy and ten controls), a test set (ten and ten), and a training set with the remaining participants. Data from participants was not shared between these three datasets.

Each model was trained in the training set and then evaluated in the validation set. The best performing wavelet type in the validation set was used to train the classifier one more time on the pooled training and validation set. Best model performance in the validation set was defined based on classification accuracy. If the accuracy was equal between models, the performance was based on - in this order - the Cohen’s kappa coefficient, area under the precision-recall curve, and or sensitivity. Finally, the best model from the validation set was evaluated in the test set, both in the same country and the test set from the other country.

Subset selections were repeated with eleven unique seeds (R seed: 100 to 600 with steps of 50, and identical between conditions) to bootstrap performance estimates in the test set. Additionally, variable importance was evaluated using the decrease in Gini score per feature.

Logistic regression was used to test for associations between binary classification success and sex, age, and data quality. A P value < 0.05 was considered significant.

## 3. Results

### 3.1 Participant characteristics

A total of 97 individuals from Guinea-Bissau and 212 from Nigeria enrolled in the study. Demographics of the participants corresponding to the data used are shown in Table 1.

**Table 1:**
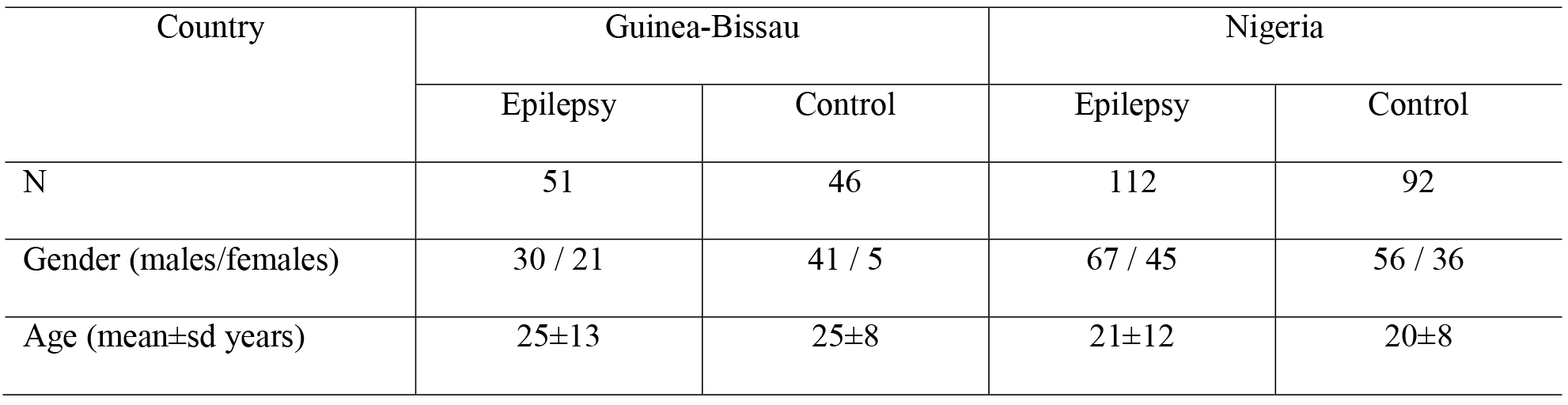
Demographics.

### 3.2 Data quality

The data acquired in Guinea-Bissau had slightly lower quality compared to the data acquired in Nigeria. The average minimum Emotiv data quality score across the channels was: 4 (very good) in 97% and 80% of the Nigerian and Guinea-Bissau data, respectively and 3 (good) in 3% and 18% of the of the Nigerian and Guinea-Bissau data respectively. No difference was found in the proportion of data with quality score 1 (very bad) or 2 (bad) between the countries. The overall quality score produced by the Emotiv headset on the sensor impedance quality was slightly lower in Guinea-Bissau (average: 305±11) in comparison to Nigeria (average 310±20).

The proportion of the remaining data for which artifacts were detected using our own algorithm was 39% and 15% in the Guinea-Bissau and Nigerian data, respectively. In Nigeria data for eight individuals were excluded as we could not select epochs due to too much signal artifacts.

### 3.3 Classification performance

We found good classification model performance in both countries (Table 2). Classification performance within the country of model training was equal or higher in Guinea-Bissau than in Nigeria. The performance of models trained with data from one country and evaluated on data from the other country was lower on all performance metrics, except sensitivity. No clear performance difference was found between the scenarios of window aggregation before or after the classification stage (Table 2).

**Table 2:** Performance in country of training and performance in other country.

| Country of training | Country of performance | Aggregate <sup>a</sup> | Sensitivity (%) | Kappa <sup>b</sup> | AUC <sup>c</sup> | Accuracy (%) |
| --- | --- | --- | --- | --- | --- | --- |
| Guinea-Bissau | Guinea-Bissau | Before | 79±3 | 0.62±0.02 | 0.85±0.02 | 81±1 |
|  |  | After | 79±3 | 0.65±0.04 | 0.87±0.02 | 83±2 |
| Nigeria | Nigeria | Before | 74±3 | 0.40±0.05 | 0.78±0.02 | 70±2 |
|  |  | After | 80±2 | 0.40±0.04 | 0.81±0.02 | 70±2 |
| Guinea-Bissau | Nigeria | Before | 85±2 | 0.17±0.03 | 0.62±0.02 | 59±2 |
|  |  | After | 90±2 | 0.10±0.04 | 0.69±0.02 | 55±2 |
| Nigeria | Guinea-Bissau | Before | 65±3 | 0.20±0.04 | 0.64±0.02 | 60±2 |
|  |  | After | 53±4 | 0.37±0.05 | 0.72±0.03 | 68±3 |

[a = whether multiple epochs were aggregated per participant before the classification stage (at feature level) or after (at probability level); b = Cohen's Kappa coefficient; c = Area under the receiver-operator curve]

### 3.4 Feature importance

Different features were important for model performance if compared between countries (Figure 3). Features based on the minimum of the standard deviation across channels were more prominent in the Guinea-Bissau models, whereas the mean standard deviation across channels in the theta band was most prominent in the Nigerian models. Differences were less pronounced when epochs were aggregated before the classification stage (Figure 3).

**Figure 3:**
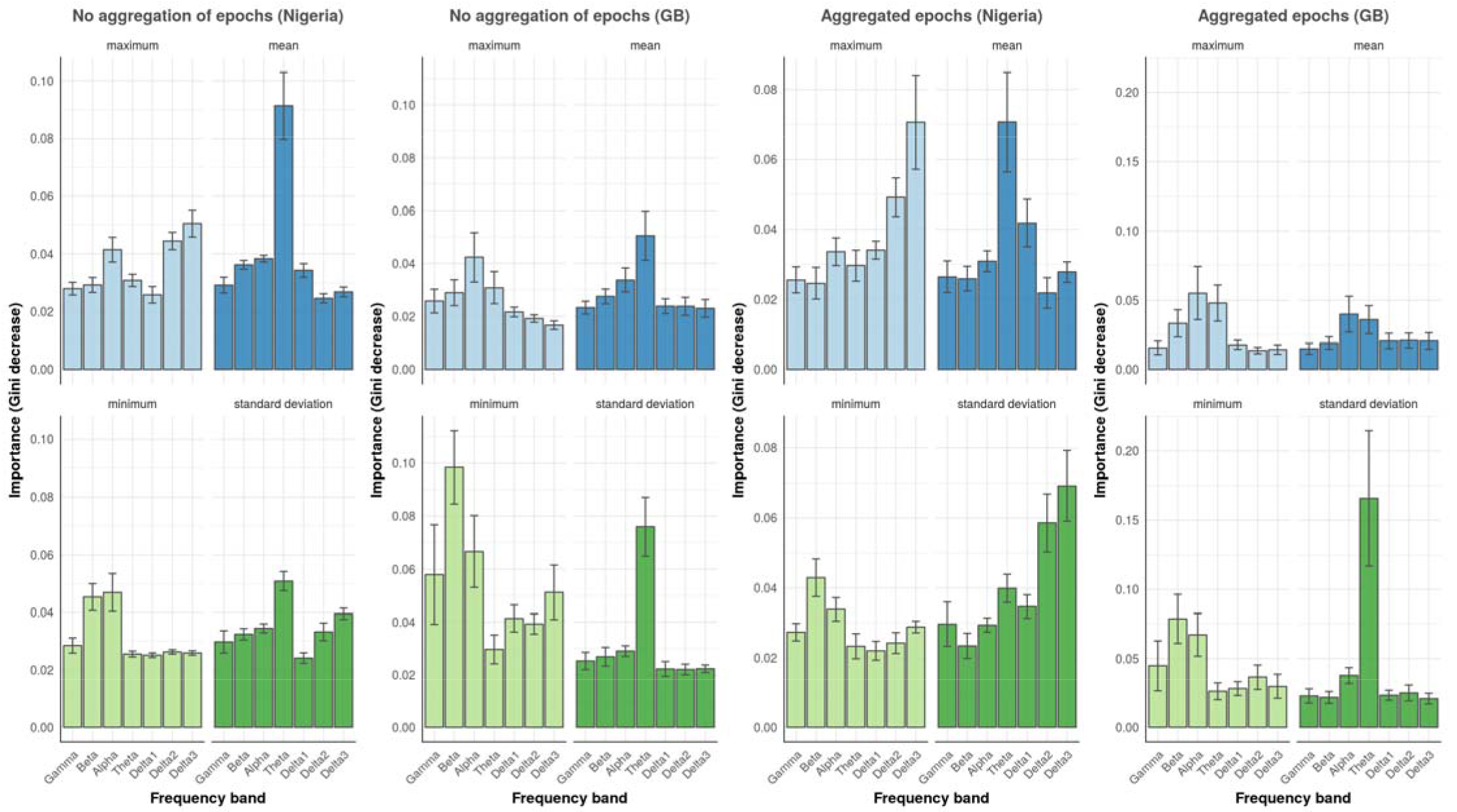
Importance of each signal feature. Values of frequency importance (i.e. decrease in Gini index) in the model performance for the Nigeria and Guinea-Bissau (GB) cohort. Values are determined for the ‘mean’, ‘minimum’, ‘maximum’ and ‘standard deviation’ of the EEG signal at individual epoch level (‘no aggregation’) or with the epoch features aggregated per participant. The frequency bands are plotted on the x-axis from high-frequency Gamma at the left to increasingly lower frequencies to the right. Errorbars represent the 95% confidence intervals.

### 3.5 Associations with classifications

No significant associations were found between classification success and sex, age, quality scores or proportion of data with artifacts.

## 4. Discussion

Our study demonstrates the potential of low cost electroencephalography to close the epilepsy diagnostic gap in difficult to reach areas in low-income and middle-income countries. The achieved within-country model performance suggests potential utility in large scale community-based screening efforts in resource-poor conditions. That is, screening in the absence of clinical experts and expensive EEG hardware. A relatively large sample size with a heterogeneous demography, acquired at multiple testing sites, outside a clinical setting, supports the robustness of our findings. The across-country performance was moderate suggesting the need for within-country optimalization.

Our approach have several advantages. The classification is based on only a few minutes of resting-state acquisition in the populations’ environment, whereas seizure calendar approaches currently in use to diagnose epilepsy in resource poor settings often take weeks to several months and requires multiple home visits by CBR workers. A second advantage is the use of easy accessible wireless EEG recordings, which do not rely on trained personal and expensive hardware. By using machine learning techniques, the data evaluation becomes less dependent on the direct availability of clinical experts. This screening tool is not intended to replace human experts. Rather, they could be used to preselect high-risk individuals for follow-up medical treatment in places where experts are lacking.

Our study has limitations. Noise levels and artifacts varied between countries. Possible sources of these differences may include various environmental factors, such as humidity, sunlight exposure, background noise, mobile phones signals, and random signal dropouts during the wireless ‘on the spot’ recordings. Further research is needed to establish the reasons for differences between locations and to develop machine learning approaches that can deal with this variation. Second, our analysis was based on channel-only signal characteristics. More information may be captured with brain-topology preserving metrics such as dynamic graphs or spectral brain images [17]. Furthermore, our current models are optimized based on classification accuracy. Given that we want to optimize the number of detected individuals with epilepsy, an optimization based on sensitivity may be more effective. However, in order to avoid a large number of false positives, we think the first focus should be on improving overall discriminative ability via the pathways as outlined above. Aggregating feature values from repeated measures within individuals before classification did not result in systematic higher performance. In theory, aggregating probabilities after the classification may fit better with a possible nonlinear relationship between feature value and epilepsy probability, while aggregation by averaging features may reduce the influence of measurement error. The current results seem to favor aggregating as this substantially speeds up the model training procedure without a clear visual drop in performance. The present study is a proof of concept and intended to guide further work. Before we can recommend clinical implementation a better generalizability and more insight into the sources of error will be needed. Finally, in the latest upgrade of the the Emotiv hardware (EPOC+) the offline data storage of raw EEG signal has been disabled and users are required to store the data in their Emotiv cloud service which requires internet connections during recording. This is not compatible with the resource poor settings used in our study. Therefore, it is important that alternative systems are developed and explored, that rely on open hard- and software, e.g. initiatives such as OpenEEG [18] and the affordable OpenBCI hardware project [19].

In conclusion, low cost brain recordings in combination with automatic time-series modeling may bring diagnostic capabilities to resource poor settings. As recently mentioned in a compelling review, the proliferation of Internet and smartphones in resource poor settings can increase health care access through so-called teleneurology [20]. Cooperation with local partners is essential for its success. A recent study has shown the potential impact of teleneurology for diagnosing epilepsy, using wireless consumer-grade EEG recordings [21]. We believe that our experience could be another step forward in this innovative field and may stimulate other contributions that may help to close the epilepsy diagnostic gap in regions where the burden of this disabling disease is highest.

## Acknowledgment

This work was supported by the Netherlands Organization for Scientific Research (NWO-VENI 016.168.038), the Dutch Brain Foundation [F2014(1)-06] and Netherlands eScience Center - Young eScientist Award 2015.

## Literature

[1] Ngugi AK, Kariuki SM, Bottomley C, Kleinschmidt I, Sander JW, Newton CR. Incidence of epilepsy: a systematic review and meta-analysis. Neurology 2011;77:1005–12. doi:10.1212/WNL.0b013e31822cfc90.

[2] McLane HC, Berkowitz AL, Patenaude BN, McKenzie ED, Wolper E, Wahlster S, et al. Availability, accessibility, and affordability of neurodiagnostic tests in 37 countries. Neurology 2015;85:1614–22. doi:10.1212/WNL.0000000000002090.

[3] Smith SJM. EEG in the diagnosis, classification, and management of patients with epilepsy. J Neurol Neurosurg Psychiatry 2005;76 Suppl 2:ii2–7. doi:10.1136/jnnp.2005.069245.

[4] Smith SJM. EEG in neurological conditions other than epilepsy: when does it help, what does it add? J Neurol Neurosurg Psychiatry 2005;76 Suppl 2:ii8–12. doi:10.1136/jnnp.2005.068486.

[5] Pillai J, Sperling MR. Interictal EEG and the diagnosis of epilepsy. Epilepsia 2006;47 Suppl 1:14–22. doi:10.1111/j.1528-1167.2006.00654.x.

[6] van Diessen E, van der Maas F, Cabral V, Otte WM. Community-based rehabilitation offers cost-effective epilepsy treatment in rural Guinea-Bissau. Epilepsy Behav 2018;79:23–5. doi:10.1016/j.yebeh.2017.11.009.

[7] David Hairston W, Whitaker KW, Ries AJ, Vettel JM, Cortney Bradford J, Kerick SE, et al. Usability of four commercially-oriented EEG systems. J Neural Eng 2014;11:46018. doi:10.1088/1741-2560/11/4/046018.

[8] Indiradevi KP, Elias E, Sathidevi PS, Dinesh Nayak S, Radhakrishnan K, Mooij AH, et al. Incidence of epilepsy: a systematic review and meta-analysis. Sensors (Basel) 2011;11:57–67. doi:10.3390/s110707110.

[9] Grummett TS, Leibbrandt RE, Lewis TW, DeLosAngeles D, Powers DMW, Willoughby JO, et al. Measurement of neural signals from inexpensive, wireless and dry EEG systems. Physiol Meas 2015;36:1469–84. doi:10.1088/0967-3334/36/7/1469.

[10] Emotiv Inc. Emotiv Systems. Emotiv - brain computer interface technology. n.d. http://www.emotiv.com (accessed April 14, 2017).

[11] Ngugi AK, Bottomley C, Chengo E, Kombe MZ, Kazungu M, Bauni E, et al. The validation of a three-stage screening methodology for detecting active convulsive epilepsy in population-based studies in health and demographic surveillance systems. Emerg Themes Epidemiol 2012;9:8. doi:10.1186/1742-7622-9-8.

[12] Janjarasjitt S. Performance of epileptic single-channel scalp EEG classifications using single wavelet-based features. Australas Phys Eng Sci Med 2017;40:57–67. doi:10.1007/s13246-016-0520-4.

[13] Mooij AH, Frauscher B, Amiri M, Otte WM, Gotman J. Differentiating epileptic from non-epileptic high frequency intracerebral EEG signals with measures of wavelet entropy. Clin Neurophysiol 2016;127:3529–36. doi:10.1016/j.clinph.2016.09.011.

[14] Indiradevi KP, Elias E, Sathidevi PS, Dinesh Nayak S, Radhakrishnan K. A multi-level wavelet approach for automatic detection of epileptic spikes in the electroencephalogram. Comput Biol Med 2008;38:805–16. doi:10.1016/j.compbiomed.2008.04.010.

[15] van Hees VT. EEG-epilepsy-diagnosis 2017. doi:10.5281/zenodo.1043938.

[16] Breiman L. Random Forests. Mach Learn 2001;45:5–32. doi:10.1023/A:1010933404324.

[17] van Diessen E, Otte WM, Braun KPJ, Stam CJ, Jansen FE. Improved diagnosis in children with partial epilepsy using a multivariable prediction model based on EEG network characteristics. PLoS One 2013;8:e59764. doi:10.1371/journal.pone.0059764.

[18] OpenEEG. OpenEEG n.d. http://openeeg.sourceforge.net/doc/ (accessed June 12, 2017).

[19] No Title n.d. http://openbci.com/ (accessed November 9, 2017).

[20] Dorsey ER, Glidden AM, Holloway MR, Birbeck GL, Schwamm LH. Teleneurology and mobile technologies: the future of neurological care. Nat Rev Neurol 2018;14:285–97. doi:10.1038/nrneurol.2018.31.

[21] McKenzie ED, Lim ASP, Leung ECW, Cole AJ, Lam AD, Eloyan A, et al. Validation of a smartphone-based EEG among people with epilepsy: A prospective study. Sci Rep 2017;7:45567. doi:10.1038/srep45567.

